# Echolocation-like model of directed cell migration within growing tissues

**DOI:** 10.1101/2022.05.13.491825

**Authors:** Tricia Y. J. Loo, Harsha Mahabaleshwar, Tom Carney, Timothy E. Saunders

## Abstract

During development and regeneration, cells migrate to specific locations within growing tissues. These cells can respond to both biochemical signals and mechanical cues, resulting in directed migration. Such migration is often highly stereotypic. Yet, how cells respond to migratory signals in a robust manner within a growing domain remains an open problem. Here, we propose a model of directed migration in growing tissues motivated by echolocation. The migrating cells generate a signaling gradient that induces a response signal from the moving system boundary. This response signal mediates cellular adhesion to the surrounding matrix and hence modulates the cell migration. We find that such a mechanism can align a series of cells at stable positions within growing systems and can effectively scale to system size. Finally, we discuss the relevance of such a model to fibroblast migration and location within the developing zebrafish caudal fin, which may be regulated by opposing signaling gradients of Slit-Robo pathway components.

**Significance Statement:** How do cells reliably migrate within growing environments? Here, we show that cells can take advantage of an echolocation-like process, whereby they induce a response from the tissue boundary. As they approach the boundary, the response signal strengthens and brings the cell to a fixed position from the boundary. This simple system may be applicable to fibroblast migration in the fin.

---

**D**uring development, there is substantial tissue growth as organs develop (1, 2). Subsequently, tissues take on a broad variety of shape and forms (3–6), with cells precisely located within the tissue (7–9). Theoretical modelling approaches have been important in understanding how mechanical forces between cells drive tissue morphogenesis (10–14). Yet, despite advances in understanding growth at a tissue scale, it remains an open problem as to how cells find specific locations within growing tissues.

The question of how cells migrate during development has received substantial attention (15). Chemotaxis, whereby cells follow specific extracellular chemical signalling gradients (16, 17), plays an important role in directing cell migration in many systems, including *Drosophila* oogenesis (18, 19), immune cell response (20, 21) and single cell migration (22, 23). Cells can also follow gradients in tissue stiffness, which is termed durotaxis (24–31). Further, flow generated gradients of mechanical stiffness (32) can play a role in processes such as cancer progression (33). The role of integrins (34) and other mechansensitive molecules (35) and the extracellular matrix (ECM) (36) have been identified as key in facilitating directed cell migration (37–39). Confined migration within dense tissues can also be influenced by mechanical properties of the nucleus (40) and can require cell division, which alters the local mechanical environment (41). In summary, directed cell migration can take many forms and often requires the interplay of biochemical and biomechanical inputs.

Collectively, cells can generate their own signals to direct migration (42, 43). In this case, a specific subpopulation becomes defined as leader cells. These cells can act as a sink, degrading the chemoattractant, and hence amplifying the local chemotactic gradient (42). Self-generated gradients appear to play a role in migration of the zebrafish lateral line (44–46) and in *Drosophila* border cell migration (47). Models of such self-generated gradients have shown how collective behavior between the leader and follower cells is essential in robust migration (48, 49). During neural crest migration (50), both pushing and pulling mechanisms between cells are important in directing cell movement (51). Neural crest cells undergo contact inhibition (52) and display collective behavior (53–55) whereby only a few cells are needed to provide direction (51); even a leading edge of migrating cells is not necessarily required (56).

Due to the multiple scales on which collective cell migration occurs (from biochemical signals, to cells, to whole tissue), modelling has been important in conceptualising mechanisms for robust migration (57–60). Methods include cell-scale approaches (61), agent-based simulations (62) and continuum models (63). Modelling has provided important insights, such as the crossover from single to collective cell behavior (64) and the role of interactions between migrating cells and the surrounding matrix (65).

Here, we focus on systems that are themselves growing significantly during the process of cell migration. How do cells precisely and robustly position within tissues that are themselves growing? We propose a model akin to a form of echolocation: (1) cells generate a signal that is received at the system boundary; (2) at the boundary a return signal is generated; and (3) the return signal modulates cell-tissue adhesion, thereby controlling cell migration. We show that this model can locate cells precisely within growing tissues, including scaling to system size. Further, it can be extended to multiple cells. Our approach does not include direct cell-cell interaction, but instead the effects of multicellularity are mediated via the tissue boundary. We apply this approach to describing fibroblast migration in the zebrafish fin fold and show that it gives results consistent with experimental observations.

## Theoretical Model

To begin, we consider a cell at position *x*_*c*_ in a one-dimensional system of length *L*. The cell generates a chemical signal, denoted by concentration *S*, at rate *J*_*S*_. The signal *S* then diffuses through the system, with diffusion coefficient *D*_*S*_, and effectively degrades at rate *µ*_*S*_ (Fig. 1A) (note, that the effective degradation can incorporate processes such as protein trapping or loss from the system). The signal concentration profile is then given by:

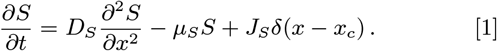

**Fig. 1.**
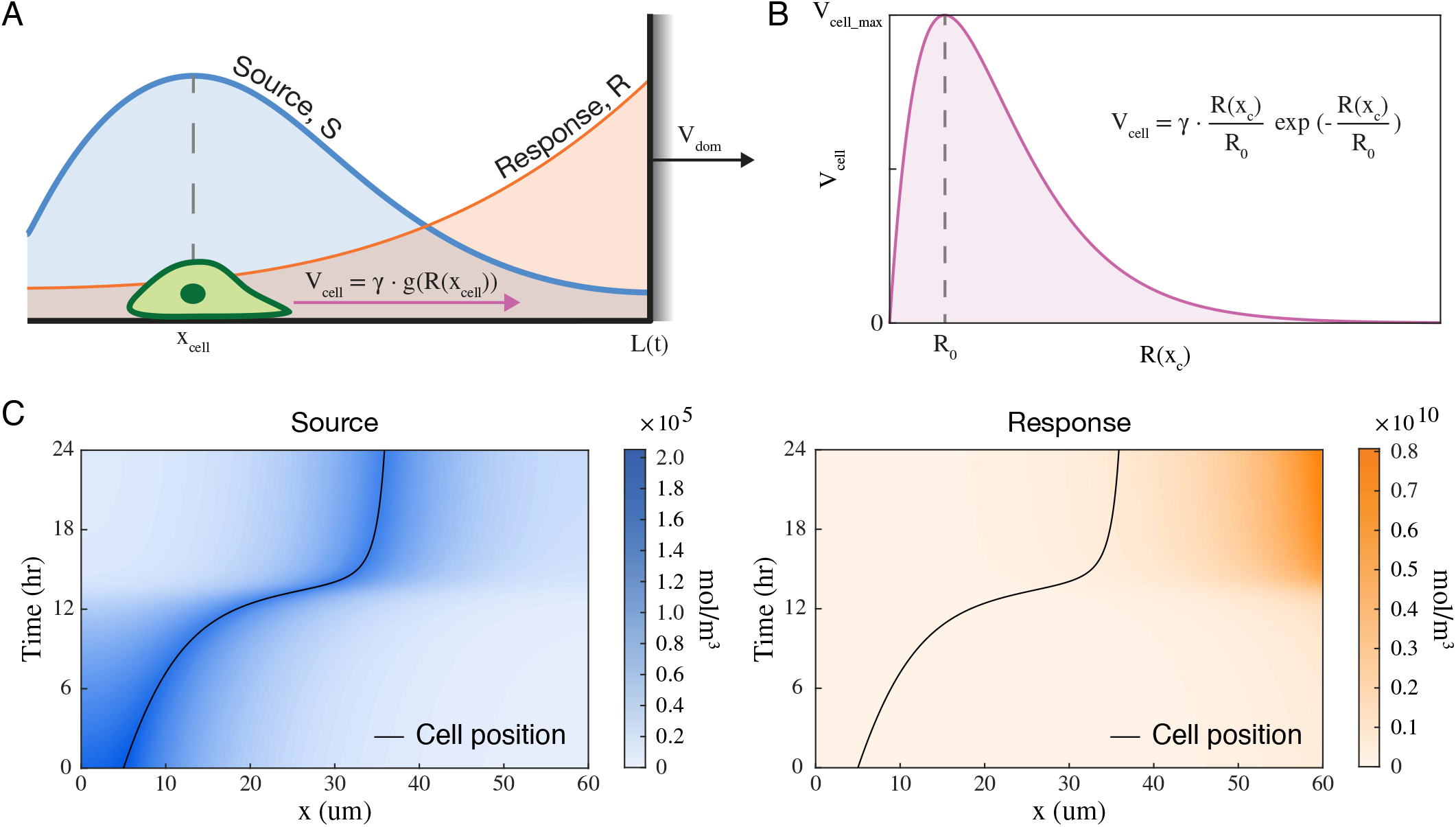
An ‘echolocation’ model for cell migration. (A) Illustration of the model for a single cell migrating in one-dimension. A cell migrates in a system of length, *L*(*t*), that grows at rate *V*_*dom*_. The cell generates a source signal, *S*(*x, t*), that elicits a response signal, *R*(*x, t*), from the system boundary at *x* = *L*(*t*), proportional to *S*(*L*(*t*), *t*). The cell speed, *V*_*cell*_, is a function of the response signal that it detects at *R*(*x*_*c*_, *t*). (B) Cell migration speed is modelled with the phenomenological function *g*(*y*) = *γ · ye*^*−y*^, where *y* = *R*(*x*_*c*_)*/R*_0_. (C) Kymograph of cell position and signal concentrations for a single cell migrating on a fixed one-dimensional domain (*V*_*dom*_ = 0*µm*) of length *L* = 60*µm*, with initial position *x*_*c*_ = 5*µm* and reaction parameters given in Table 1.

At the boundary, the signal generates a return signal, denoted by concentration *R* (Fig. 1A). The flux of *R, J*_*R*_, is assumed to be a function of *S*(*L*). The return signal diffuses with diffusion coefficient *D*_*R*_ and effectively degrades at rate *µ*_*R*_:

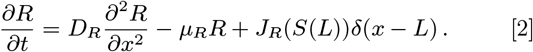

From Eq. 1 and 2, we can define effective length scales 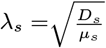 and 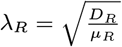.

The system itself grows. For simplicity, we take growth to be constant at a rate *V*_*dom*_,

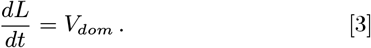

Finally, we assume that the cell responds to the return signal *R*(*x*_*c*_), which determines its rate of motion:

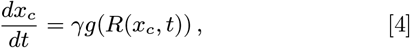

where *g*(*y* = *R/R*_0_) represents the haptotactic response of the cell to the concentration of *R. R*_0_ is a characteristic concentration determining the sensitivity of the response of the cell to *R*(*x*_*c*_)· *γ* is a constant with dimensions of velocity. The precise form of *g*(*y*) is system dependent, but has the general form of *g*(*y* ≪ 1) ≈ 0 and *g*(*y* ≫1) ≈ 0, with a single maximum for positive *y* and *g*(*y*) *>* 0 for *y >* 0. At small *y*, this corresponds to the cell having weak contact with the surrounding substrate (like walking on ice). At large *y*, this corresponds to the cell adhering very strongly to the surrounding substrate (like walking through mud). At intermediate values, the cell migrates at a maximum velocity. Here, we use the phenomenological function *g*(*y*) = *ye*^*−y*^ (Fig. 1B). Our results are not dependent on the specific form of this function so long as it obeys the above general conditions.

In the simulations, we are interested in the behavior of *x*_*c*_ (position of the cell), and its dependence on the system’s growth. The biochemical signals likely propagate rapidly compared with the speed of cell migration, meaning that we can consider Eq. 1 and 2 to be in steady-state compared to Eq. 4. Hence,

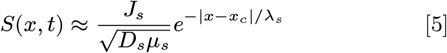

This assumes the cell is a point source. Within the simulations, it is straightforward to implement the source coming from the cell boundaries. From Eq. 5, we can find the concentration of *S* at the boundary,

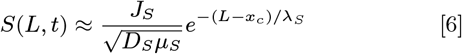

Subsequently, assuming that *J*_*R*_(*S*(*L, t*)) = *j*_*R*_*S*(*L, t*) (*i*.*e*. the production rate scales linearly with the signal concentration) we have

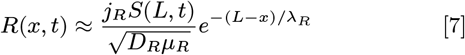

which leads to the response concentration at the cell given by

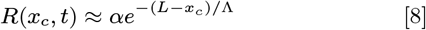

where 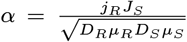 and 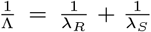 describes the effective signal production and effective decay length respectively. For more than one cell and with growing boundaries, it becomes more challenging to make analytical progress. Therefore, we utilize simulations for exploring the system behavior in more complex scenarios.

## Results

### Biphasic response of cell velocity to signal allows for boundary detection

We first explored how a single cell migrates in one-dimension towards a fixed edge. The cell generates its own signal, which induces a response from the boundary (see Methods). In Fig. 1C, we show a single cell reaching its final position in the tissue. As the cell approaches the boundary, *R*(*x*_*c*_) gradually increases, causing the cell to slow to a stop due to increased adhesion.

At low velocities, any directed motion is nullified by randomness inherent in the system; we can assume that at some threshold value *g*_*c*_ such that *g*(*R*(*x*_*c*_)*/R*_0_) ≤ *g*_*c*_, *dx*_*c*_*/dt* = 0 on average. The velocity response crosses the threshold value at either extremes of *R*. Therefore, if the cell starts too far from the leading edge, it will not initiate migration. Likewise, if it starts close to the leading edge, then it will not move relative to the edge throughout growth. Here, we always initiate the system in a condition such that cell migration is possible.

We define a concentration of *R* such that the cell comes to its final position, *x*_*c,stop*_ when *R*(*x*_*c,stop*_) is greater than or equal to *R*_*stop*_. Therefore, we can find the position at which the cell ceases to migrate relative to the leading edge:

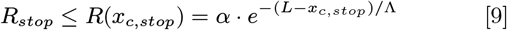

*L − x*_*c,stop*_ gives the stopping distance of the cell from the boundary and it depends on the reaction-diffusion parameters of the biochemical signals involved. Since the cell slows to a stop as it approaches the boundary, the stopping distance is given by its maximum value:

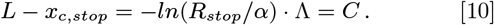

For each set of reaction parameters, there is a particular stopping distance; here, we are only interested in parameter regimes where *x*_*c,stop*_ exists within the system length, *i*.*e*.*C* ∈ [0,*L*].

### Scaling cell migration

We can rearrange Eq. 10 to get the relative cell stopping position of the cell:

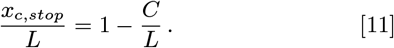

The relative stopping position is reciprocal to the domain length; this allows the cell stopping distance to exhibit a degree of scaling as the domain length changes (Fig. 2A). We numerically verified this by running the simulations with different domain lengths *L*. At very small system lengths, the cells receive a high *R* from the boundary immediately, and hence they fail to migrate far. At larger system sizes, the eventual relative stopping distance does not vary significantly (for our parameters in Table 1, this corresponds to systems ≳75*µm*). Fig. 2B gives the normalized cell tracks for each simulation. Though the cell trajectories are quite different in different system sizes, changes in domain length only affected the amount of time needed for the migrating cell to come to a stop, with the migrating cells eventually converging on a small range of relative stopping positions.

**Fig. 2.**
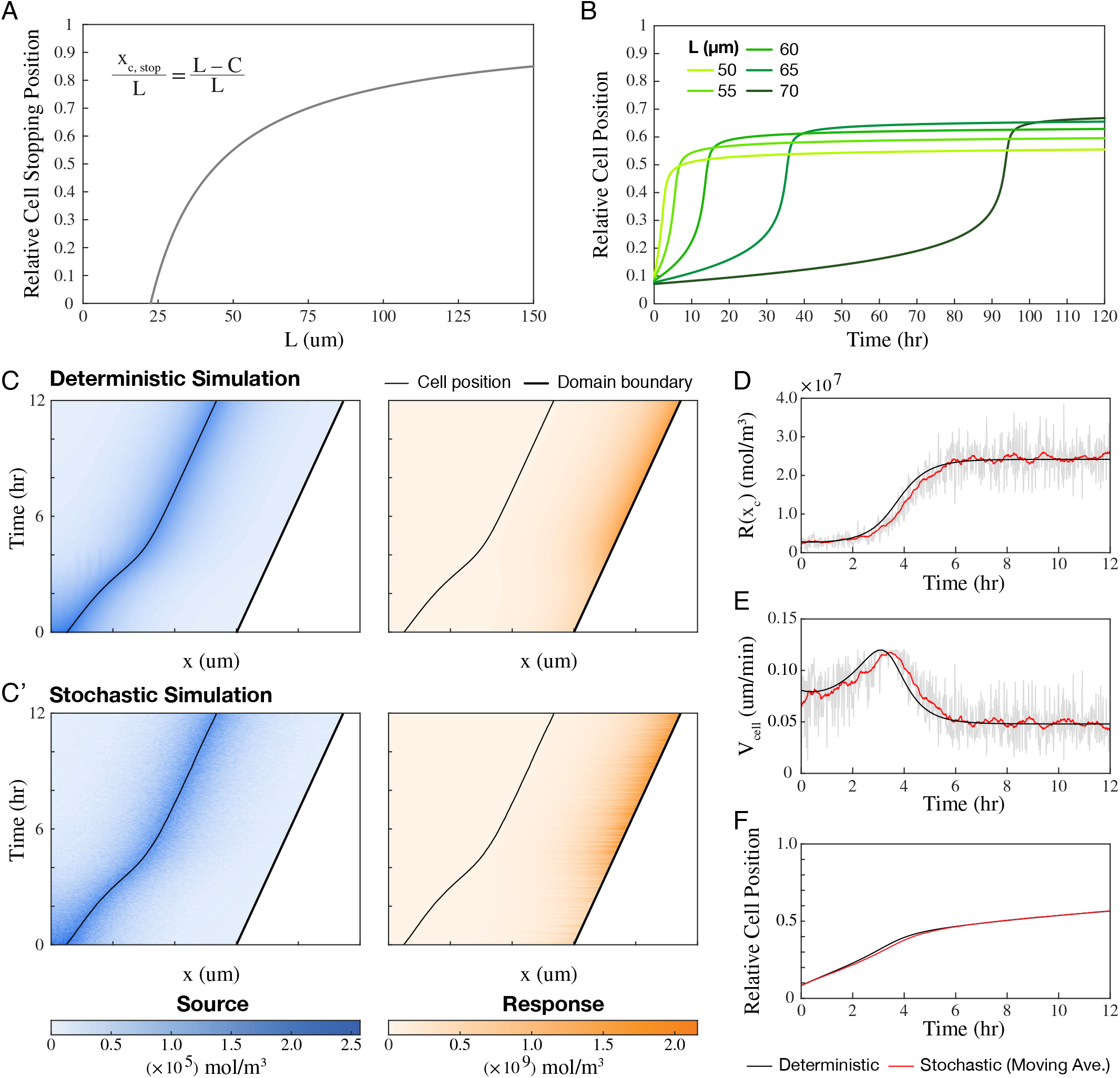
Biphasic cell response gives rise to robust boundary detection and scaling. (A-B) The relative cell stopping position is robust to large changes in the system length. (A) Analytical solution of relative cell stopping position vs. system length for parameters in Table 1. *R*_*stop*_ is chosen such that *R*(*x*_*c*_) ≥*R*_*stop*_ gives *V*_*cell*_ ≤1 *×*10^*−*6^*µm/s*. (B) Numerical simulations of a single cell migrating in systems of fixed lengths *L* = 50, 55, 60, 65, and 70*µm, V*_*dom*_ = 0*µm/min*. (C-F) Growth is introduced to the system, *V*_*dom*_ = 0.048*µm/min*, and *R*_0_ = 8 *×*10^6^*mol/m*^3^; to investigate robustness, we solve the system using both the deterministic numerical simulation and a stochastic particle-based simulation. (C, C’) Kymographs of cell position with concentration of source signal (left) and response signal (right). (D-F) Plots of *R*(*x*_*c*_) (D), *V*_*cell*_ (E) and relative cell position *x*_*c*_*/L*(*t*) (F), over time as the system grows. Black: deterministic simulation; gray and red (F): stochastic simulation; red (D-E): moving average from stochastic simulation.

**Table 1.** Parameters used in simulations unless stated otherwise.

| Parameters | $D_S, D_R$ | $\mu_S, \mu_R$ | $J_S, j_R$ | $R_0$ | $\gamma$ |
| --- | --- | --- | --- | --- | --- |
| Values | $10\mu m^2 s^{-1}$ | $0.1 s^{-1}$ | $0.3 mol m^{-2} s^{-1}, 0.3 s^{-1}$ | $10^8 mol m^{-3}$ | $0.33 \mu m min^{-1}$ |

We extended this result to explore domains that change in length over time. When domain growth is implemented in the simulation, as shown in Fig. 2C-F, the cell can track the leading edge of the system and shows distinct phases of acceleration and then constant velocity while tracking the leading edge. In the simulation, at around *t* = 6hr, the migrating cell reaches its stopping position relative to the boundary, then maintains a fixed distance from the growing boundary (Fig. 2C). This is because the cell can only move forward when the domain grows and *R*(*x*_*c*_) falls back below the stopping concentration *R*_*stop*_. This can be seen in Fig. 2D-E, where *R*(*x*_*c*_) approaches a steady value such that the cell migrates at the same rate as boundary growth. When the cell has reached its final position relative to the boundary, this is stable despite continued system growth (Fig. 2F).

We tested the robustness of these results to intrinsic fluctuations in the cell motion and the reaction kinetics (Methods). Though single simulation tracks were noisy (gray curves in Fig. 2D-E), they still displayed the distinct shifts in cell velocity and scaled stopping distance (Fig. 2F). The average behavior was similar to the deterministic solution, suggesting that this mechanism of echolocation can robustly guide cells to specific locations within growing tissues.

Overall, we have shown that an echolocation-like process can direct cells to specific locations within growing domains, and this process is robust to fluctuations in the system components.

### Collective migration: Neighbor detection through feedback signaling

Having described the behavior of single cells in growing domains, we next asked how do multiple cells respond to the echolocation-like interaction with the boundary? Under self-directed signaling, the amount of *R* generated at the organising boundary increases as the migrating cell approaches the boundary. When more than one cell is present, each cell will have different velocity profiles across the length of the system. As the leading cell approaches the boundary, the trailing cell receives a greater amount of *R* signal than the leading cell did when it was in the same position. Hence, the leading and trailing cells reach their final stopping points at different positions within the domain (Fig. 3A-B).

**Fig. 3.**
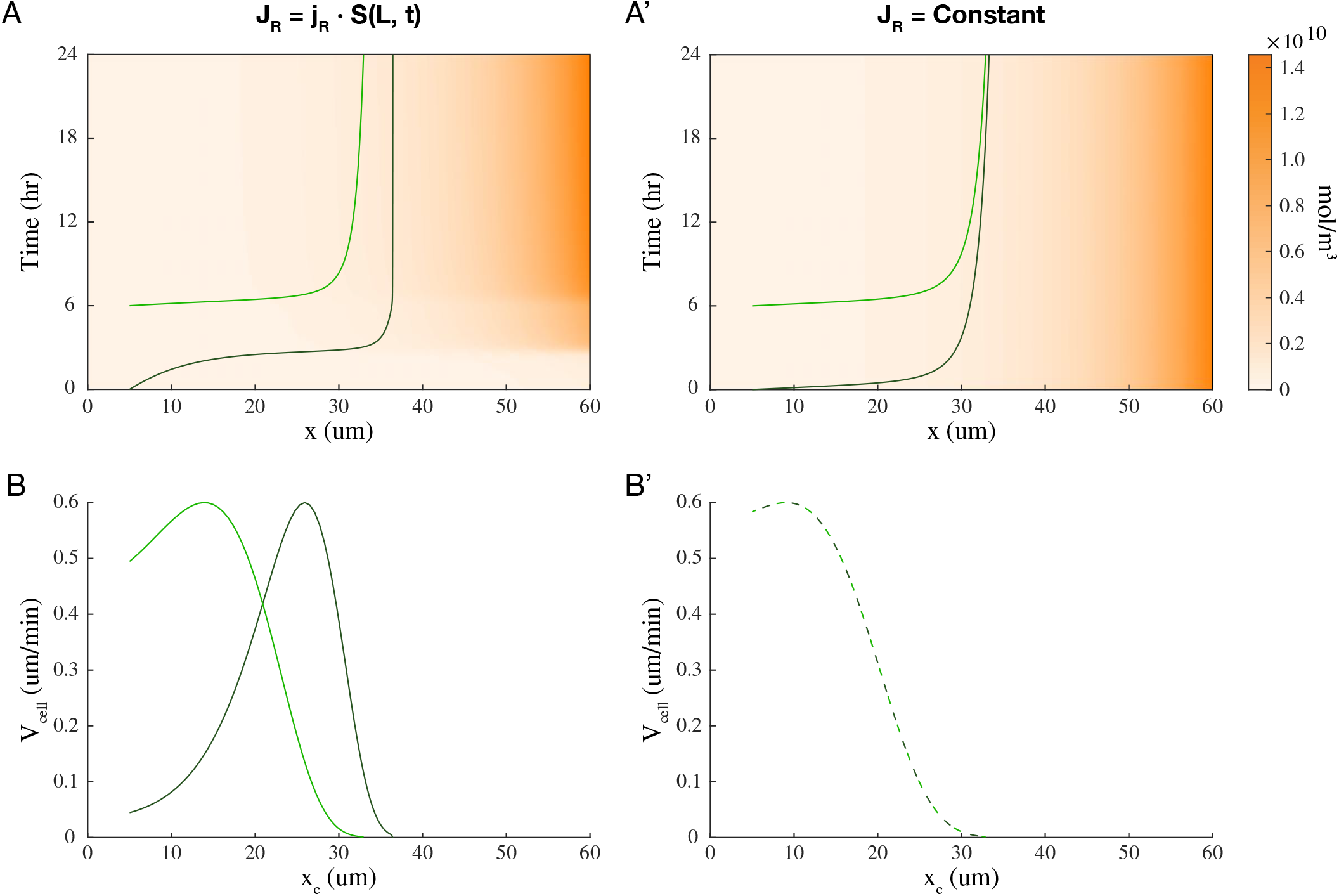
Self-directed signaling allows for neighbor detection. Two cells migrating in a fixed one-dimensional system of length *L* = 60*µm*, in response to either a self-generated signal (left) or a constant signal (right; *J*_*R*_ = 1.4 *×* 10^4^*mol/*(*m*^3^ *· s*)), with *γ* = 1.6*µm/min*. (A, A’) Kymographs of R concentration and each cell track; (B, B’) Plot of *V*_*cell*_ against *x*_*cell*_ position for each cell in the system.

In comparison, when the organising boundary produces a constant signal, the concentration profile of *R* does not change in time, and all migrating cells will have the same velocity profile across the length of the system. Both leading and trailing cells will have the same stopping position and will tend to converge (Fig. 3A’ and 3B’). Echolocation-like behavior can enable cells to position separately within a domain without need for processes such as contact inhibition.

### Domain growth and cell initial position shapes cell behavior

How sensitive is our model to the chosen parameters? In this section, we test the robustness of the model to parameters. The boundary tracking behavior seen above does not occur in all circumstances in the simulations. In Fig. 4A-B, we can see system growth outpacing cell migration. The boundary receives less of the source signal and in return produces less of the response, resulting in the cell coming to a stop far from the boundary. How do the reaction parameters affect cell migration behaviour and in what parameter regime are cells able to track the boundary throughout boundary growth?

**Fig. 4.**
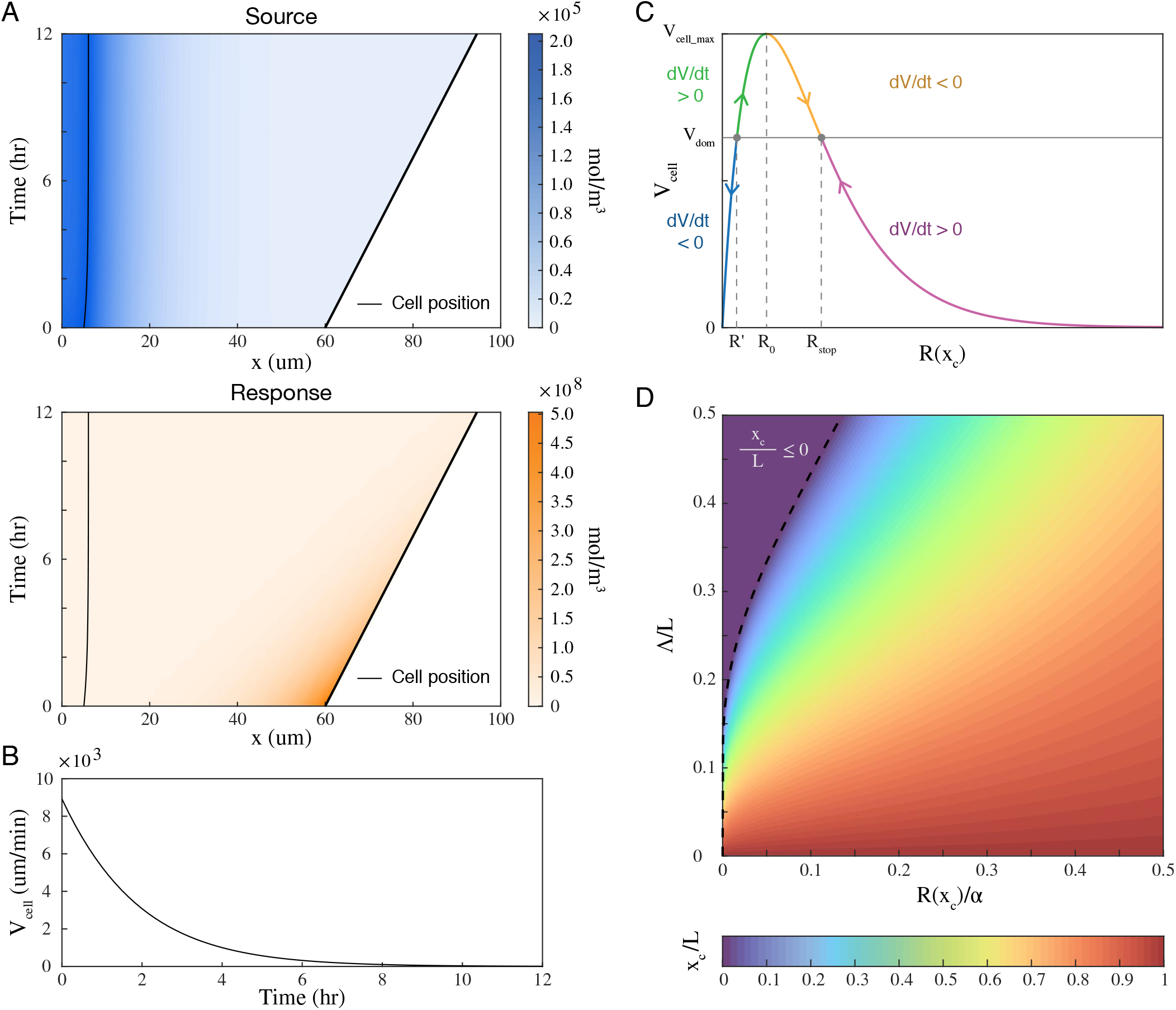
Cell initial position, domain growth rate and *R*_0_ value determines cell behavior. (A-B) A single cell migrating in a growing domain with change in system length, *V*_*dom*_ = 0.048*µm/min*, and starting position *x* = 5*µm*. Kymograph of cell position, signal and response concentration (A) and cell velocity over time (B). (C) The acceleration profile of the cell in response to *R* is defined by the domain growth rate *V*_*cell*_ = *V*_*dom*_ and maximum cell response at *R*(*X*_*cell*_) = *R*_0_. Since *R* varies monotonically with *x*, the initial position of the cell and its initial *R*(*x*_*cell*_) will determine if the cell will migrate to *x*_*stop*_ or be left behind by the growing boundary. (D) Parameter space of relative cell positions *x*_*cell*_*/L* as a function of the relative effective decay length Λ*/L* and generalized reaction parameter *R*(*x*_*cell*_)*/α*. As the effective decay length Λ increases, the other parameters are more constrained to ensure cell position remains bounded by the system length [0, *L*].

Our cell migration response function is modelled by:

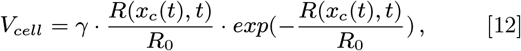

from which we can get the instantaneous acceleration of the cell,

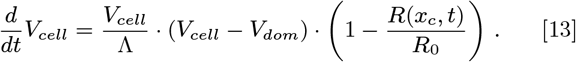

The acceleration of the cell is dependent on two factors: (1) whether the domain growth rate is greater than the cell migration rate, *V*_*dom*_ *> V*_*cell*_; and (2) whether *R*_*x,cell*_ is greater than the critical response threshold, *R*_0_ (Fig. 4C). There are only two stable points in the system at *V*_*cell*_ = 0 and *V*_*cell*_ = *V*_*dom*_. There is additionally an unstable equilibrium point to the left of *R*_0_, such that *V*_*dom*_ = *V*_*cell*_, which we will call *R*,. If *R*(*x*_*c*_) *< R*^*′*^, then the cell will decelerate and stop moving altogether (blue regime in Fig. 4C), but if *R*(*x*_*c*_) *> R*^*′*^, then the cell will eventually move at the same rate as system growth, tracking the boundary.

We first consider the effects of domain growth rate. If the domain length is fixed, *V*_*dom*_ = 0, the entire system falls within the green and yellow regimes in Fig. 4C, so all cells will be able to detect the boundary and eventually slow to a stop at *x*_*cell,stop*_. If *V*_*dom*_ *> V*_*cell,max*_, then system growth will always outpace cell migration and the cells will stop because they have lost the ability to sense the boundary (blue and pink regimes in Fig. 4C). However, if the domain growth is at some intermediate rate between 0 and *V cell, max*, then the cell behavior is determined by whether *R*(*x*_*c*_) is greater or less than *R*^*′*^.

We next look at the effects of the *R*_*x,cell*_. Since *R* is produced at the rightmost boundary in our setup (*x* = *L*), *R* increases monotonically along the system length, and serves as a proxy for cell position. Any cell which has an initial position left of position *x*^*′*^, where *R*(*x*^*′*^) = *R*^*′*^, will be unable to catch up to system growth, whereas cells that begin to the right of this *x*^*′*^ will stop at some constant distance from the boundary. Where this critical position is located depends on the number of cells already present in the system and their current positions. It is possible that as leading cells migrate towards the boundary, sufficient *R* production induces the entourage of trailing cells to begin migration, when otherwise they may have failed to migrate if acting individually.

To test the sensitivity of our model to the particular parameters chosen, we next explored the parameter space determining the cell final position, within the constraint that *x*_*c*_*/L* ∈ [0, 1]. Rearranging the Eq. 8, we can express the relative cell position as

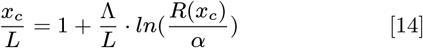

Since all parameters are expressed in positive values, Λ*/L* is also positive, and *R*(*x*_*c*_)*/α* is bounded by [0, 1]. We explore this parameter space in Fig. 4D, with Λ*/L* ≤ 0.5, such that the domain length is at least twice that of the effective signal decay length. If *x*_*c,stop*_*/L <* 0, the cell never initiates migration, whereas if *x*^*′*^*/L >* 1, all cells will be left behind by the growing system. As the effective decay length Λ of the system increases, the available space for *R*(*x*_*c*_) = *R*^*′*^ or *R*_*stop*_ to exist decreases. However, when Λ is small, the positions of *x*^*′*^*/L* and *x*_*stop*_*/L* will decrease in distance and come closer to the boundary at *x*_*c*_*/L* = 1.

### Fibroblast Migration in Zebrafish Fin Fold

We have developed a model of how long-ranged biochemical interactions between a cell and the boundary can guide cell migration, both singularly and collectively. To explore whether this has *in vivo* relevance, we focused on the developing zebrafish embryo fin. During early fin formation, fibroblast cells migrate distally from the proximal mesoderm region towards the apical ectodermal ridge (AER), moving along the inner wall of the epidermis, Fig. 5A. Details of the experimental system can be found in Ref. (66). Given the very narrow width of the fin, cell migration within this system is effectively two-dimensional.

**Fig. 5.**
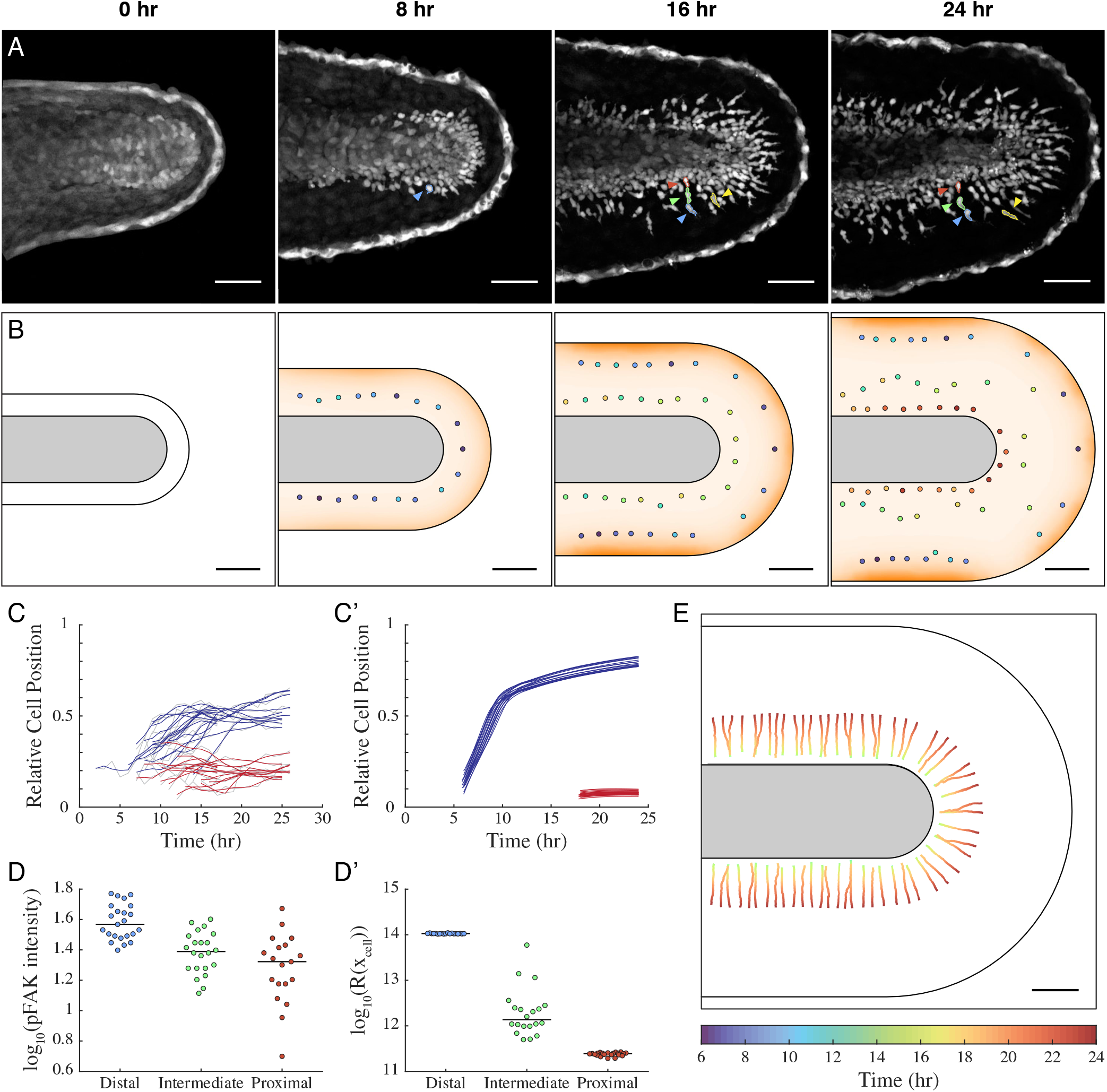
Modelling mesenchymal cell migration in zebrafish median fin fold. (A) Development of zebrafish median fin fold (time since 24hpf). Three representative cell tracks depict migration behavior of distal (blue), intermediate (green, yellow) and proximal (red) cells. (B) Simulation of cell migration in two dimensions with Response signal concentration; *R*_0_ = 3.5 *×* 10^13^*molm*^*−*3^. (C, C’) Normalised tracks of distal (blue) and proximal (red) cells from embryos (C) and from simulations (C’). (D, D’) Log-scaled jitter plots of pFAK intensity in embryos (D) (replotted from (66), Fig. EV5) and *R* signals in simulation (D’). Immunofluorescent staining shows a gradient of focal adhesion activity in cells across the fin, corresponding with the gradient of R signalling expected from simulations. (E) Cell tracks from a single simulation in which each cell acts as a sink for *R. R*_0_ = 5 *×* 10^3^*molm*^*−*3^. Scale bar = 50*µm*.

Genetic experiments by Mahabaleshwar et. al (66) suggest that this migration is mediated by feedback signals between the cell and the AER. Slit ligands are produced by the migrating fibroblasts, which then act via its canonical receptor Robo at the AER to induce the release of Sphingosine-1-Phosphate (S1P). S1P then may form a signaling gradient across the fin that is high distally and low at the proximal fibroblasts, *i*.*e*. with the opposite profile to the Slit gradient from the cell. Sphingosine-1-Phosphate Receptor 2 (S1PR2), which is expressed in the migrating fibroblast, is known to be involved in cell migration signaling *in vivo* (67–69). This effectively alters the local cellular adhesive environment, thereby regulating the speed of cell migration.

We implemented this network within our simulation environment. We represented the fin as a two-dimensional environment with cells migrating from the proximal mesoderm region toward the AER, Fig. 5B. The results recapitulate the *in vivo* system, where the most distal cells display boundary tracking behavior, and the cells that are far from the boundary do not migrate very far. We performed single cell tracking to dissect the cellular dynamics (Methods). In both experiment (Fig. 5C) and simulation (Fig. 5C’) the first cells to migrate (blue lines) initially migrated quickly towards the AER. As they approached the AER they slowed down and then approximately tracked the tissue boundary growth. This is due to increased cell adhesion. Cells that entered the system later (red lines) stayed largely stationary and did not track with the growing tissue boundary.

A key prediction of the model is that the local adhesive environment should change with respect to distance to the AER. In our model, the adhesive environment is related to the level of *R*(*X*_*cell*_). In Ref. (66) the levels of the adhesion adhesion molecule pFAK (70) were analyzed across the fin fold. Comparing this data (replotted in Fig. 5D) with *R*(*X*_*cell*_) (Fig. 5D’), we see that the levels of adhesion and *R* are correlated. Note, we do not expect to see a precise correlation as the response signal *R* is interpreted by the cell in a likely nonlinear manner. To test whether this relationship is functionally affecting cell migration requires more controlled perturbation of the local adhesive environment (*e*.*g*. by optogenetics) during cell migration.

In the simulations, neighbor-neighbor interactions differ slightly from the one-dimensional case, as trailing cells are no longer spatially constrained behind the leading cells. The trailing cells can thus catch up to join the ranks in the spaces between leading cells. Though we saw signatures of such behavior in the experimental data (yellow cell in Fig. 5A), it is hard to quantify this due to challenges in defining the trailing cells.

Overall, our model can be applied to cell migration within the developing zebrafish fin fold. It is consistent with both cell scale movements and also local changes in the mechanical environment.

## Discussion

Here, we have developed a model of directed cell migration within growing domains that bears similarity with echolocation mechanisms. By cells effectively communicating with themselves and each other via the system boundary, cells can position reliably within a growing domain, both in terms of maintaining separation between cells and their distance from the boundary. We have taken a continuous approach to describing both the molecular, cellular and tissue scale processes. We have shown that this mechanism is robust to reasonable levels of intrinsic noise and parameter variations. It appears that the feedback between the cells and the system edge act to buffer variability. Our model results are largely consistent with observed behavior of migrating fibroblasts in the developing zebrafish fin fold (Fig. 5). Our model predicts that the local adhesive environment changes with distance from the AER, and it will be interesting to test this observation further *vivo*. More quantitative measurements on cell migration in the fin fold will aid comparison to experiment as interpreting cell collective motion data is subject to error (71).

Modelling of cell migration behavior can depend on the level of model approximation (*e*.*g*. discrete vs continuous approaches, (72)) and it will be interesting to try different approaches within our echolocation framework. Though we do not see substantial changes in the dynamics when the cells are stochastically modeled (Fig. 2D-F), a more detailed analysis may reveal important differences. In particular, noise may determine cell behavior near the critical response thresholds (Fig. 4C).

A fascinating behavior we observed in our simulations of the tail fin is that the leading cells tended to cluster towards each other rather than spread out. As a migrating cell approaches the boundary, it generates more *R* at the nearest points of the boundary and attracts other cells to move in a similar direction. Experimentally, we do not see this behavior in the tail fins. This discrepancy may be explained by other cell-cell interactions, such as BMP-mediated short-ranged contact inhibition (73, 74), that is not modelled within our framework. In our model, cells respond to the return signal *R*, but do not alter the local *R* concentration; *i*.*e*. the cells do not act as sinks for *R*. If the cells act as sinks, then (particularly at high cell densities) it is possible that the cells themselves can modulate the level of *R* available in the system. Similar to chemotaxis (75), this may sharpen the local gradient of the response signal. This could then have an impact on the polarity and adhesion to substrate of the following cells. We performed an initial simulation with the cells acting as sinks (Fig. 5E). Cells which appeared later in the simulation caught up with the more distal cells but then spread out to avoid crowding (branching seen in Fig. 5E). It will be interesting to explore such behavior further theoretically and to test whether such processes occur *in vivo*.

Further analysis can explore other growing systems that also have concurrent cell migration to see whether such “echolocation” plays a role in driving robust cell movement.

## Materials and Methods

### Simulation Framework

We solved our above model using numerical simulations in different geometries. We first explored the model in 1D. At each time step, we created a mesh of the system with the current cell position and current domain size, then solved for the steady state solutions of Eq. 1 and 2. We extracted the signal concentration at the current cell position, calculated the cell velocity and updated the cell position to generate a new mesh for the next time step. We assumed that growth takes place only at the boundary *x* = *L*, rather than as an expansion of the entire domain. As such, the cell position *x*_*c*_ is not affected by system growth.

To test the robustness of the model, we also solved the 1D model stochastically. At each time step *dt*, particles representing *S* and *R* are randomly added to and removed from the system with a Poisson probability and rate parameters determined by *J*_*S*_ · *dt, j*_*R*_ · *S*(*L*) · *dt, µ*_*S*_ · *S* · *dt* and *µ*_*R*_ · *R · dt* respectively. The particles then move a random distance drawn from a Normal distribution with mean *µ* = 0, and variance *σ*^2^ = 2 ·*D*_*S*_ · *dt* and *σ*^2^ = 2 · *D*_*R*_ · *dt* respectively. We obtain the concentration of *R*(*x*_*c*_) by taking the mean concentration of *R* over *x* = *x*_*c*_ ± 2.5*µm*. From this, we find the cell migration rate, and update the cell position and domain size for the next iteration.

Unless otherwise specified, all simulations use the reaction parameters given in Table 1.

The numerical simulations in 2D follows the same regime as in the 1D case, with the exception that the direction of cell migration is determined by the spatial gradient of *R*, while the magnitude of the velocity vector is calculated from the response curve (Eq. 4). We additionally explored a sink condition in which the cells degraded the response signal R at the rate of *−*0.001*m*^2^*s*^*−*1^ · *R*(*x*_*c*_, *t*).

### Imaging and Analysis

We compare the simulation results to fibroblast cell migration in the developing zebrafish fin fold. Cell lines, movies and pFAK intensity measurements were obtained as described in Ref. (66). Selected cells were manually segmented and tracked on Fiji. The relative cell position of each cell was measured as 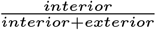, where *interior* is the shortest distance between the cell centroid and the proximal mesoderm, and where the *exterior* 455 was the shortest distance between the cell centroid and the apical ectodermal ridge.

## ACKNOWLEDGMENTS

This work was funded by Singapore Ministry of Education MOE Tier 3 grant (MOE2016-T3-1-005) (all authors) and University of Warwick startup funding for T.E.S.

